# Histone demethylase KDM5 regulates cardiomyocyte maturation by promoting fatty acid oxidation, oxidative phosphorylation, and myofibrillar organization

**DOI:** 10.1101/2023.04.11.535169

**Authors:** Manisha Deogharia, Akanksha Agrawal, Miusi Shi, Abhinav K Jain, Kevin J. McHugh, Francisco Altamirano, A J Marian, Priyatansh Gurha

**Author notes:** Address for Correspondence Priyatansh Gurha Center for Cardiovascular Genetics Brown Foundation Institute of Molecular Medicine The University of Texas Health Science Center, 6770 Bertner Street, C950G.

## Abstract

**Rationale:** Human pluripotent stem cell-derived CMs (iPSC-CMs) are a valuable tool for disease modeling, cell therapy and to reconstruct the CM maturation process and identify, characterize factors that regulate maturation. The transition from immature fetal to adult CM entails coordinated regulation of the mature gene programming, which is characterized by the induction of myofilament and OXPHOS gene expression among others. Recent studies in *Drosophila*, *C. elegans,* and C2C12 myoblast cell lines have implicated the histone H3K4me3 demethylase KDM5 and its homologs, as a potential regulator of developmental gene program and mitochondrial function. We speculated that KDM5 may potentiate the maturation of iPSC-CMs by targeting a conserved epigenetic program that encompass mitochondrial OXPHOS and other CM specific maturation genes.

**Objectives:** The purpose of this study is to determine the role of KDM5 in iPSC-CM maturation.

**Methods and Results:** Immunoblot analysis revealed that KDM5A, B, and C expression was progressively downregulated in postnatal cardiomyocytes and absent in adult hearts and CMs. Additionally, KDM5 proteins were found to be persistently expressed in iPSC-CMs up to 60 days after the onset of myogenic differentiation, consistent with the immaturity of these cells. Inhibition of KDM5 by KDM5-C70 -a pan-KDM5 inhibitor-resulted in differential regulation of 2,372 genes including upregulation of Fatty acid oxidation (FAO), OXPHOS, and myogenic gene programs in iPSC-CMs. Likewise, genome-wide profiling of H3K4me3 binding sites by the CUT&RUN assay revealed enriched H3K4me3 peaks at the promoter regions of FAO, OXPHOS, and sarcomere genes. Consistent with the chromatin and gene expression data, KDM5 inhibition led to increased expression of multiple sarcomere proteins, enhanced myofibrillar organization and improved calcium handling. Furthermore, inhibition of KDM5 increased H3K4me3 deposits at the promoter region of the *ESRRA* gene, which is known to regulate OXPHOS and cardiomyocyte maturation, and resulted in its increased RNA and protein levels. Finally, KDM5 inhibition increased baseline, peak, and spare oxygen consumption rates in iPSC-CMs.

**Conclusions:** KDM5 regulates the maturation of iPSC-CMs by epigenetically regulating the expression of ESRRA, OXPHOS, FAO, and sarcomere genes and enhancing myofibril organization and mitochondrial function.

## INTRODUCTION

The replicative capacity of human cardiomyocytes (CMs) declines progressively during cardiac development as these cells gradually differentiate toward mature cells. Consequently, the fully differentiated adult CMs possess minimal if any residual replicative capacity ^1, 2^. The loss of replicative capacity of mature CM in the adult heart prevents regeneration and replacement of CMs following myocardial infarction; a leading cause of heart failure ^3-5^.

The transition of proliferative and immature CMs to mature and fully differentiated cells involves a series of morphological, metabolic, and functional changes ^6, 7^. The changes are driven by a switch from a fetal-like to an adult gene expression profile encompassing genes involved in the myofibrillar organization, mitochondrial oxidative phosphorylation (OXPHOS), Fatty acid oxidation (FAO), and an isoform switch in several sarcomere proteins. These changes are also responsible for a shift in metabolic substrate utilization from glucose and lactate to free fatty acids as an energy source. All of these changes eventually lead to improved OXPHOS and sarcomere alignment and increased contractility, among others ^8^.

The recent discoveries that allow efficient differentiation of human pluripotent stem cells (PSCs) into the CM lineage upon induction of the cardiogenic program have increased interest in the utility of these cells to replace the damaged CMs. However, the immaturity of the induced PSC-derived CMs remains a major bottleneck in their functional utility and has therefore shifted interest to interventions that can enhance iPSC-CM maturation ^9-12^.

Current approaches to induce maturation of iPSC-CMs includes engineered micropatterned platforms, synthetic cardiac tissue, electrical stimulation, prolonged culture time, and biochemical modification (transcription factor and signaling component alteration), which have exhibited varying degrees of effectiveness ^11^. Recent studies have linked chromatin organization and epigenetic modifications such as DNA methylation and histone marks, including the trimethylation of histone H3 at lysine residue 4 (H3K4me3) to CM development and differentiation ^6, 7, 13-15^. In the heart, activating histone modifications, such as H3K4me3, and H3K27ac are enriched in gene promoters associated with CM maturation^14^. H3K4me3 is the main target of histone lysine demethylases KDM5A and B, which we recently found to be activated in CMs in a mouse model of lamin-associated cardiomyopathy^16^. Members of the KDM5, comprised of KDM5A (chromosome 12), KDM5B (chromosome 1), KDM5C (X chromosome), and KDM5D (Y chromosome) are a subfamily of JmjC KDMs, which act as transcriptional corepressors by catalyzing the removal of methyl marks from mono-di- an trimethylated H3K4 ^17^. Recent studies in *Drosophila*, *C. elegans* and C2C12 myoblast cell lines have revealed a role of KDM5 and its homologs, as a potential regulator of developmental gene program and OXPHOS gene expression^18, 19^. Furthermore, we proposed that the KDM5 family of proteins by reducing H3K4m3 levels at the promoter regions of the genes encoding mitochondrial OXPHOS and cell cycle regulators suppress their expressions in the heart ^13, 16^. Given that the metabolic regulation of iPSC-CMs is a determinant of maturation, we hypothesized that KDM5 family members are involved in iPSC-CM maturation by targeting conserved and cell type-specific gene programs that encompass mitochondrial OXPHOS among others.

In the current study, we elucidated the role of KDM5 in cardiomyocyte maturation. Inhibition of KDM5 switched the transcriptome toward a more mature CM gene expression profile and increased the expression of transcription factor ESSRA and genes involved in OXPHOS, sarcomere formation, and fatty acid metabolism. The changes were associated with increased OXPHOS and enhanced myofibril formation, indicative of a shift toward maturation of the iPSC-CMs.

## METHODS

### Isolation of neonatal cardiomyocytes

Neonatal cardiomyocytes were isolated from 2-day-old mouse neonates, as published. ^20^ In brief, the heart was removed and immediately transferred to a petri dish containing an ice-cold ADS buffer pH 7.3 (6.8 g NaCl, 0.4 g KCl, 1 g dextrose, 1.5 g NaH_2_PO4, 0.1 g MgSO_4_, 4.76 g HEPES). After the removal of blood, the hearts were transferred to a petri dish containing a digestion buffer consisting of 3650 units of collagenase II, 30 mg pancreatin, and 50 ml of ADS buffer. The hearts were cut into small pieces in the digestion buffer and transferred to a small flask fitted with a stir bar. The flask was placed on a hot plate at (37°C, 200-400 rpm) and the hearts were digested for 4 min. The supernatant was discarded and the tissue was digested in 5 ml of the digestion buffer for 10 min. After collecting the supernatant, 1 ml of Bovine calf serum was added to stop the reaction. This process was repeated 5-6 times and the supernatant was collected in each round. The supernatants were pooled and passed through 100 µm cell strainers. After filtration, the cells were centrifuged at 1000 rpm for 4 min. The pellets were resuspended in 12 ml of cardiomyocyte (CM) medium and plated on a culture dish at 37°C for 1 hour to allow the non-CMs to adhere to the plastic. After 1 hour, media with CMs were collected from the Petri dish and centrifuged at 1000 rpm for 3 min. The isolated myocytes were suspended in urea buffer for immunoblotting assay.

### Isolation of adult cardiomyocytes

Adult cardiomyocytes were isolated from P14, P28, and P56 mice as published ^13, 16, 20-23^. Briefly, the heart was excised from the anesthetized mouse, mounted on a Langendroff perfusion system, and perfused with collagenase digestion buffer (2.4 mg/ml, Worthington cat. No LS004176) at a flow rate of 4 ml/min. Upon completion of the digestion, (generally 10-12 min) the blood vessels and atria were excised, and the ventricles were minced in a stop buffer containing 10% calf serum, 12.5µM CaCl_2_, and 2mM ATP. The cell suspension was filtered through a 100µm cell strainer and centrifuged at 20 g for 4 minutes to precipitate cardiomyocytes. The isolated cells were treated with a gradient of CaCl2 concentration ranging from 100µM to 900µM in the stopping buffer. The isolated myocytes were suspended in a urea buffer for immunoblotting.

### Maintenance and differentiation of human induced pluripotent stem cells (iPSC)

Commercially available human iPSC lines derived from a female donor (Gibco Episomal hiPSC, Gibco, catalog # A18945) and from a male donor (ChiPSC22, Cellartis catalog # Y00320, Takara) were maintained in mTeSR1 medium (STEMCELL Technologies, catalog #85850) in a six-well dish coated with 300µg/ml Matrigel (Corning, catalog #354230). Cells were checked for mycoplasma contamination using the Mycoplasma detection kit (Southern Biotech, catalog #13100-01) according to the manufacturer’s protocol. Cells were split (1:9) at 70-80% confluency using EZ-LiFT Stem Cell Passage Reagent and maintained for 24 hours in mTESR1 medium plus 10µM rock inhibitor Y-27632 (STEMCELL Technologies, cat. 72308). Cells were grown in mTESR1 and the media was changed every 24 hours until cells were 75-85% confluent. Most experiments were performed in Gibco Episomal hiPSCs (henceforth hiPSC). To initiate differentiation, the iPSCs were treated (day 0) with CHIR (9µM for hiPSC) in RPMI/B27 without insulin (RPMI/B27-) for 48 hours. Exactly 48 hours later (Day 2), the medium was switched to IWP2 (7.5µM) in RPMI/B27- and on day 4 to RPMI/B27- only. Every 48 hours thereafter the medium was changed to RPMI/B27 with insulin (RPMI/B27+) ^24-27^. Post-differentiation (day 9) wells that showed consistent beating throughout were used for subsequent experiments.

### Inhibition of KDM5

After differentiation, iPSC-CMs were lifted from the plate with TrypLE Express (Gibco, #12605028), filtered through a 100µM filter, and placed in RPMI/B27+ medium with a ROCK inhibitor (10µM) and 10% knockout serum for 24 h. After 24 h, cells were treated with a series of concentrations of pan-KDM5 inhibitor KDM5-C70 ^28-30^ (xcessbio, M60192) and 100µM BSA-conjugated palmitate (1:3), and the medium was changed every 3 days for 15 days (D15– D30). The experimental groups to determine KDM5 inhibition efficiency were as follows: 1) iPSC-CMs, 2) iPSC-CMs plus DMSO (0.01%) 3) iPSC-CMs + KDM5-C70 (0.1µM), 4) iPSC-CMs + KDM5-C70 (0.5µM), 5) iPSC-CMs + KDM5-C70 (2µM) and iPSC-CMs + KDM5-C70 (10µM). Optimal KDM5 inhibition, as indicated by H3K4me3 accumulation, was found at a concentration of 0.5µM and therefore all subsequent experiments were performed using this concentration of KDM5-C70.

### Immunoblotting

Immunoblotting (IB) was performed as published^13, 16, 23, 31-34^. Cells were lysed using Pierce RIPA buffer (Pierce, Catalog#89901) with 0.5% SDS or urea-thiourea buffer (containing 7M urea, 2M thiourea, 1% SDS and 50mM Tris-Cl pH 6.8) plus protease and phosphatase inhibitor and were sonicated for 3 cycles (30s ON and 30s OFF each cycle) using Bioruptor. The extracts were centrifuged at 14000 rpm for 15’ at 4°C or RT for urea-thiourea buffer. Aliquots of 10-15µg of extracts were loaded onto SDS-PAGE and electrophoresed. Proteins were transferred to a nitrocellulose membrane overnight at 4°C. The primary and secondary antibodies are listed in Supplementary Table 2.

### Structural integrity and sarcomere organization

Cells after treatment for 15 days with KDM5-C70 were removed by using a 1:1 mixture of TrpLE and Accutase (Invitrogen, A11105-01) and plated on glass coverslips coated with Matrigel (300µg/ml). Cells were allowed to recover for two days and fixed and stained with anti–ACTN2 and TNNT2 antibodies. Quantitative assessments of circularity, cell area, and perimeter were done by selecting the cell boundaries and using ImageJ software.

### Preparation of PDMS-coated coverslips

Sylgard 184 PDMS (Dow Corning, Midland, MI, USA) was prepared by combining 9:1 (w/w) base:curing agent, mixing thoroughly, and centrifuging for 3 min at 300 g to remove bubbles. An aliquot of 10 µL of the uncured PDMS solution was pipetted on each glass coverslip (12 mm circle) and then each PDMS-coated coverslip was flipped over and placed PDMS side down onto fluorinated ethylene propylene (FEP) sheet. Coverslips were then placed under a vacuum, to further remove bubbles, and then placed in a 70 °C oven for 30 min to cure. Coverslips with adherent PDMS were then carefully peeled from the FEP sheet, flipped over, and cured in an oven at 120 °C for at least 2 h. PDMS-coated coverslips were then sterilized in 70% ethanol, washed with phosphate-buffered saline, air dried, and plasma treated before ECM coating.

### Quantitative real-time PCR

Transcript levels of selected genes were quantitated by qPCR, using gene-specific primer and SYBR Green master mix, and data were normalized to GAPDH or Vinculin (VCL). The sequences of qPCR primers are included in Supplementary Table 1.

### RNA sequencing

RNA-seq was performed as published previously^13, 16, 20, 32, 34^. Briefly, the RNeasy Mini Kit was used to isolate total RNA. The samples with an RNA integrity number (RIN) greater than 9.0 were used for sequencing. The ribosomal RNA was removed and the RNA was then processed for strand-specific sequencing. Samples were sequenced on an Illumina Nextseq500 using 75 bp paired-end sequencing. STAR aligner was used to map the reads to the human Hg19 genome and Feature Counts program was used to quantify the reads^35, 36^. Genes with a read count of at least one read per million in at least three samples were used for further analysis. The R/Bioconductor package limma-voom ^37^ was used to calculate the fold change and adjusted p-value (q-value). Genes with a fold change of 1.5 and q< 0.01 in KDM5-C70 versus cells and DMSO were considered as differentially expressed.

### Cleavage Under Targets and Release Using Nuclease (CUT&RUN) assay

Cut-and-run was performed at the MD Anderson Cancer Center Epigenomics Profiling Core according to published protocols ^38, 39^. Briefly, ∼500,000 untreated and KDM5-C70 treated iPSC-CMs were harvested, washed, and attached to activated Concanavalin A-coated magnetic beads, then permeabilized with wash buffer containing 0.05% digitonin. Upon H3K4me3 antibody binding and washing, the DNA was released, purified and the libraries were created using NEB Ultra II Library prep kit following manufacturer’s instructions. Size distribution of the libraries was determined using the Agilent 4200 TapeStation and libraries were multiplexed to achieve uniform representation. Libraries were sequenced using Illumina Nextseq 500 instrument to obtain 15-20 million reads per sample. Processing of the sequencing data was performed using Pluto Bioscience software(https://pluto.bio/), which used Bowtie2 for mapping to the human genome build Hg38 and SEACR for peak calling^40^. Consensus peaks from all samples were determined with minimum overlap set of 2 and differential expression of peak counts was performed using Diffbind (summit size 500bp) on local galaxy instance. Further processing of the data was done on the local galaxy instance that entails mapping the differentially expressed to the corresponding genes using Homer software (annotate peak) and data visualization using the Deep Tools package to identify genome enrichment and functional annotations. The genomic distribution of the differential peaks was also performed by PAVIS (https://manticore.niehs.nih.gov/pavis2/).

### Pathway analysis

Gene set enrichment analysis (GSEA) was performed on normalized counts using the gene sets curated from the Molecular Signature Database (MSigDB) 3.0. Significance was assessed by analyzing signal-to-noise ratio and gene permutations based on 1,000 permutations. The data were ordered based on the enrichment score for the gene set with a q-value < 0.05. The Upstream Regulator Analysis module of Ingenuity Pathway Analysis software (IPA, QIAGEN Redwood City) was used to identify likely transcriptional regulators. DEGs were used for this analysis.

### Mito Stress assay

After 15 days of treatment (with either DMSO or KDM5 inhibitor), cells were removed upon treatment with a 1:1 mixture of Accutase and TrypLE. At least 40,000 cells/well were plated in Matrigel-coated 96-well dishes in RPMI/B27+ medium containing 100µM Palmitate, 10% knockout serum, and 10µM of Y-27632 (ROCKi). After 24 h of incubation, the same medium as above was added but without ROCKi, and Mito stress assays using an Agilent Seahorse machine were performed according to the manufacturer’s protocol. Briefly, one hour before the assay, cells were washed twice with Agilent Seahorse XF RPMI Basal Medium supplemented with 1mM sodium pyruvate, 2mM glutamine, and 10mM glucose (103577-100, Agilent Technologies) and incubated in the same medium for 1 hour at 37°C in a non-CO2 incubator. Ports A, B, and C of the XF96 sensor cartridge (102601-100, Agilent Technologies) were loaded with oligomycin (1.5µM), FCCP (2µM), rotenone, and antimycin A (0.5µM), respectively. The oxygen consumption rate (OCR) was normalized to the protein concentration determined by the BCA assay. Seahorse Analytics was used to calculate Basal respiration, Spare respiratory Capacity, and Maximal respiration.

### Measurements of Intracellular Ca^2+^

Cells were replated onto Matrigel-coated 35 mm glass bottom dishes (Cellvis) and allowed to recover for 4-5 days. After that, hiPSC-CMs were loaded with Fluo-4-acetoxymethyl (AM)-ester (5 μM, Invitrogen) in Tyrode’s buffer (135 mM NaCl, 5 mM KCl, 1.8 mM CaCl_2_,1mM MgCl_2_, 5.6 mM glucose, 10 mM HEPES pH 7.4) for 25 min at RT. Cells were washed three times with Tyrode’s buffer and incubated for an additional 25 min to promote Fluo-4 AM de-esterification. Cells were paced at 0.2 Hz using a pair of platinum wires, and fluorescence was monitored using an IonOptix system equipped with a 470 nm LED, emission filter, and photomultiplier. Cells were continuously paced during the entire experiment to achieve a steady-state Ca^2+^ level. Fluo-4 fluorescence is expressed as (F-F0)/F0 x 100 where F is the fluorescence intensity corrected by background fluorescence. A total of 20 consecutive transients were averaged to measure calcium amplitude, time to peak, and exponential decay constant (tau) using IonWizard and Cytosolver (IonOptix). Exponential fitting was conducted between the time point when the signal recovered 10% of the baseline and when it returned to baseline. Data from four different batches of hiPSC-CMs were analyzed per group (10cells per batch per group). Descriptive calcium parameters were evaluated in the cells which follow pacing without major arrhythmogenic events.

### Statistical analysis

The Shapiro-Wilk test was used to determine the normality of the data distribution. Ordinary one-way ANOVA was used to analyze normally distributed data, followed by Tukey’s multiple pairwise comparison test. The Kruskal-Wallis test was used to analyze data that deviated from the normal distribution, followed by Dunn’s pairwise comparisons. GraphPad Prism 9 was used to perform statistical testing. For iPSC cells, all experiments are performed as 2-3 batches with each batch having 3-4 replicates of samples obtained from independently treated wells. A subset of gene expression changes was validated by qPCR in an independent cell line ChiPSC22 (Supplementary Figure 4C, Supplementary Figure 5C, and D). RNA-seq and CUT&RUN genomic data are available from the NCBI Sequence Read archive via the following links: https://dataview.ncbi.nlm.nih.gov/object/PRJNA950177?reviewer=rp7gf0d1a0lhnuoe82ammup70t https://dataview.ncbi.nlm.nih.gov/object/PRJNA950477?reviewer=5268iv42jvoa7lftgp3rc43ptf.

## RESULTS

### The expressions of the KDM5 family of proteins are downregulated upon the maturation of murine cardiomyocytes

Immunoblotting of myocardial protein extracts from 5-day (P5), 3-month (P90), and 12-month (P180) old mice showed a progressive decline in the expression levels of KDM5A, KDM5B, and KDM5C in the postnatal period, as these proteins were expressed in the 5-day old but were absent in 3- and 12-month-old mouse hearts (Figure 1A-B). To determine whether the decline reflects changes in the expression of these proteins in cardiomyocytes, IB was performed on protein extracts from cardiomyocytes isolated from P2, P14, P28, and P56 old mice hearts. The KDM5 proteins were found to be expressed in P2 and P14 cardiomyocytes but their levels were markedly reduced in P28 and P56 cardiomyocytes (Figure 1C-D). Thus, we surmise that KDM5 expression levels decline with the maturation of murine cardiomyocytes.

**Figure 1:**
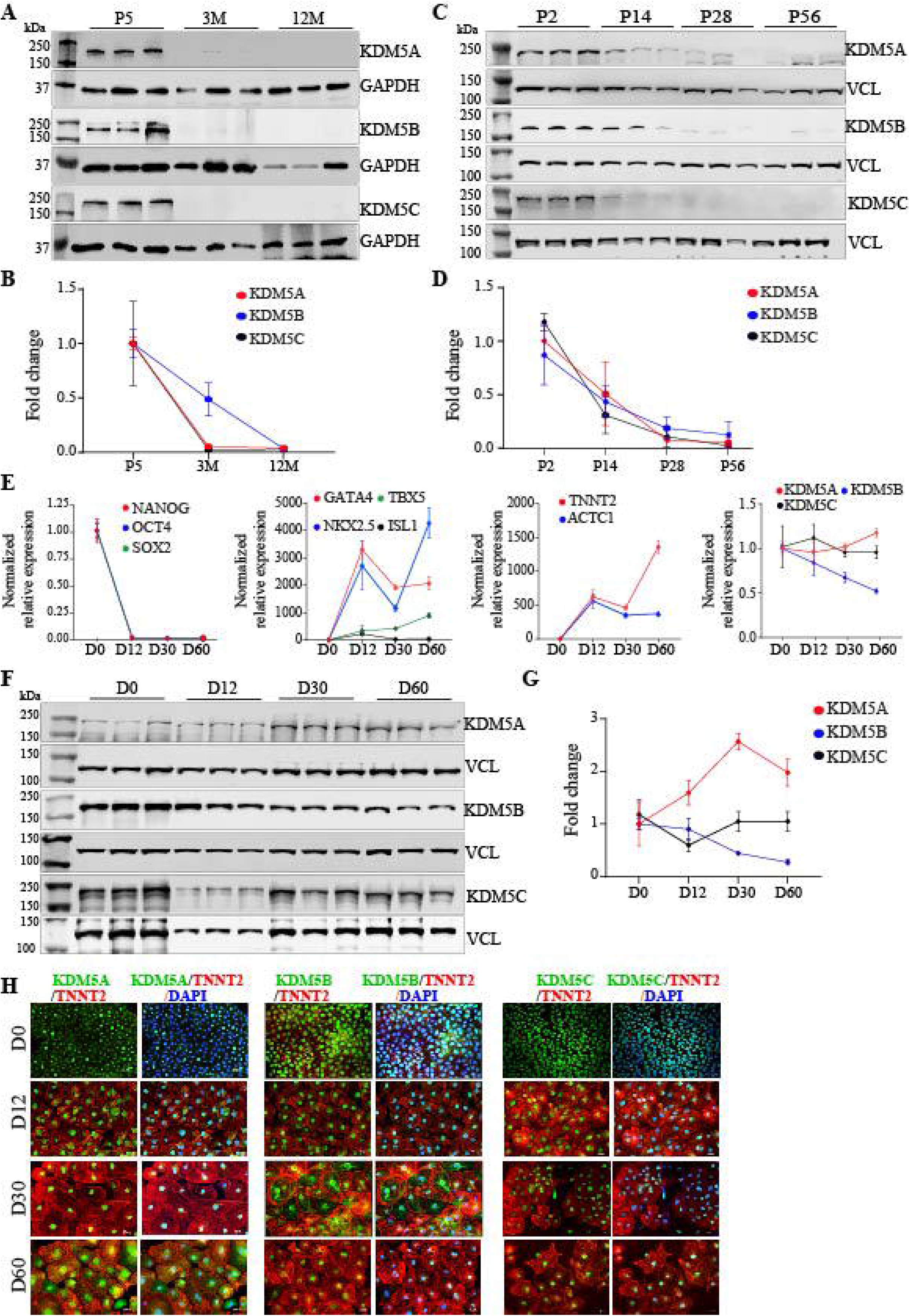
Expression of KDM5A, B and C in immature and mature cardiac myocytes. **A)** Immunoblot (IB) analysis from whole heart extracts showing the expression of KDM5A, KDM5B and KDM5C in the neonate (P5) and their absence in heart samples from adult mice. GAPDH was used as a loading control. **B)** Respective quantitative data for KDM5A, B, and C normalized to GAPDH levels from whole heart extracts. **C)** IB analysis of KDM5A, B, and C levels in mouse cardiomyocytes isolated from newborn (P2), 2-week (P14), 4-week(P28), and 8-week (P56) mouse hearts. **D)** Corresponding quantitative data for KDM5A, B, and C from cardiac myocytes normalized to Vinculin (VCL). **E)** QPCR analysis of RNA extracted at different time points of differentiation of human iPSCs toward cardiac lineage. The relative expression of pluripotent markers, cardiac transcription factors, and CM lineage markers at different stages of differentiation is shown. The expression levels of *KDM5A*, 5B and *5C* at the indicated time points are also shown. **F)** IB analysis of KDM5A, 5B, and 5C levels from extracts obtained at different time points of iPSC differentiation to CM. **G)** Quantitative data showing consistent expression of KDM5A, B, and C proteins in iPSC and at all stages of iPSC differentiation towards CM normalized to Vinculin (VCL). **H)** IF staining of cells with anti-KDM5A, B, or C (green) and anti-TNNT2 antibodies (red) and DAPI (blue) showing the expression and nuclear localization of KDM5 proteins at all stages of differentiation.

### The KDM5 family of proteins are persistently expressed in iPSC-CMs

Given that iPSC-CMs are immature cells, the KDM5 proteins would be expected to be expressed in iPSC-CMs, if they are involved in maintaining the cells in an immature state. To test this hypothesis, human iPSC-CMs were generated upon differentiation of iPSCs using a chemically defined monolayer protocol through a sequential Wnt activation and suppression protocol ^25, 26^. The efficacy of the differentiation protocol was determined by analyzing expression levels of the molecular markers of iPSC and cardiomyocytes. As shown previously ^8, 10, 11^, a stage-specific gene expression pattern was observed, which was notable for the induction of the mesodermal lineage marker T-box transcription factor T (aka Brachyury) and MESP1 at D2-6 (Supplementary Figure 1A) and suppression of expression of pluripotency markers OCT4, NANOG, and SOX2 (Figure 1E). Suppression of expression of the pluripotency markers was associated with increased expression of cardiac transcription factors NKX2-5, GATA4, and ISL1 as well as cardiac myocyte markers ACTC1 and TNNT2 (Figure 1E). IF staining of iPSC-CMs generated from this protocol showed that 88± 9.3% of cells expressed TNNT2 (Supplementary Figure 1B). The results are consistent with the differentiation stages shown by others and confirm the validity of the approach.

To determine changes in the expression of the KDM5s at different stages of differentiation, transcript levels of KDM5A, B, C, and D were analyzed by qPCR, which showed their expressions at all stages, including the early and late stages of differentiation of iPSC-CMs (Figure 1E). KDM5D gene which is located on the Y-chromosome was only expressed in male-derived iPSC cell lines and showed a similar expression pattern during iPSC differentiation (Supplementary Figure 1C). To corroborate the findings at the protein level, IB was performed, which showed persistent expression of KDM5A, B, and C proteins at several stages of differentiation of human iPSCs to iPSC-CMs (Figure 1F-G). Furthermore, immunofluorescence (IF) co-staining of iPSC-CM with anti-KDM5 and anti-TNNT2 antibodies, the latter a marker of cardiomyocytes, confirmed KDM5 expression in the iPSCs and iPSC-CMs (Figure 1H). Moreover, IF staining showed that KDM5s were predominantly located in the nucleus (Figure 1H). These findings showing persistent expression of the KDM5 proteins in immature iPSC-CMs and their absence in the late post-natal, i.e., mature murine cardiomyocytes, suggest a role for the KDM5 proteins in the maturation of cardiomyocytes.

### Inhibition of KDM5 proteins induces expression of the mature cardiomyocyte gene program

To investigate whether persistent expression of KDM5s contributes to the immaturity of iPSC-CMs, we inhibited KDM5 activity with a small molecule inhibitor KDM5-C70. KDM5-C70 is a cell-permeable derivative of KDM5-C49 that inhibits the activity of all KDM5 proteins by competing with α-ketoglutarate for binding to the active site ^28, 29^. To obtain the effective dose for inhibition, iPSC-CMs were treated with four different concentrations of the KDM5-C70 (10, 2, 0.5, and 0.1µM) for two weeks. The treatment duration was chosen to inhibit KDM5s when their expression levels were the highest (days 15-30). Cells treated with vehicle (0.01% DMSO) and untreated cells served as controls. Inhibition of KDM5 is expected to lead to the accumulation of H3K4me3 in cells, which can be quantified by IB. IB showed a dose-dependent increase in the H3K4me3 levels upon inhibition of KDM5 with KDM5-C70 (Figure 2A). The highest dose of KDM5-C70 increased the steady-state level of H3K4me3 by 11.93±1.6-fold (Figure 2A-B). The minimum dose that achieved a maximum effect was determined to be 0.5µM, which caused a 9.9±1.6-fold increase in H3K4me3 levels compared to DMSO-treated cells. No discernible changes in the levels of other histone methylations such as H3K27me3 and H3K9me3 were observed after 7 days of treatment of iPSC-CMs with 0.5 µM of KDM5-C70 (Supplementary Figures 2A and 2B). Finally, IF staining showed nuclear accumulation of H3K4me3 in the treated group (0.5µM KDM5-C70) compared to the DMSO or non-treated group (Figure 2C). Because KDM5-C70 at a dose of 0.5µM was sufficient to suppress KDM5 activity without causing non-specific effects, this dose was chosen for the subsequent experiments.

**Figure 2:**
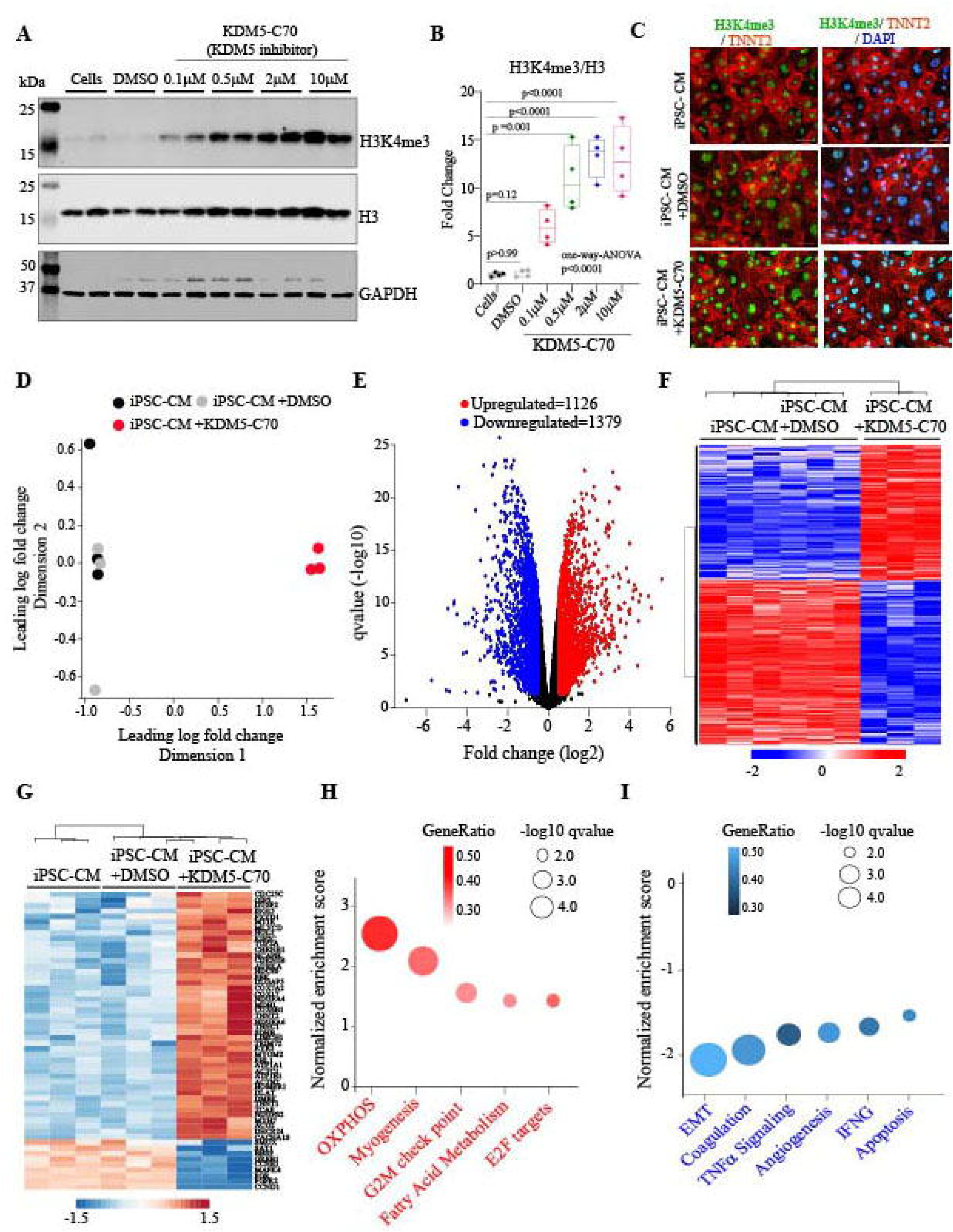
Effect of KDM5 inhibition on iPSC-CM gene expression. **A)** Immunoblot showing H3K4me3 levels after treatment of iPSC-CMs with different concentrations of the KDM5 inhibitor (KDM5-C70). Cells treated with vehicle (0.01% DMSO) and untreated cells served as controls. **B)** Changes in the level of H3K4me3 relative to H3 for the data shown in A. **C)** IF staining using anti-H3K4me3 (Green) and anti-TNNT2 (red) antibodies. **D)** Multidimensional scaling plot of RNA-Seq data showing the separation of iPSC-CMs and DMSO from the KDM5-C70 treated samples. **E)** Volcano plots obtained from RNA-seq data showing differentially expressed genes and levels of significance in KDM5-C70 compared to untreated control iPSC-CMs. The differentially upregulated genes are shown in red, while those that are downregulated are shown in blue and those that remain unchanged are shown in black. **F)** Heatmap and hierarchical clustering of differentially expressed genes in the indicated groups. **G)** Heat plot showing the expression pattern and differential expression status of KDM5A, and B predicted targets after KDM5-C70 treatment. **H–I)** Gene Set Enrichment Analysis (GSEA) after KDM5-C70 treatment showing the activated **(H)** or inhibited **(I)** hallmark gene signature. The Y-axis in the graphs represents the Normalized Enrichment Score (NES), the color intensity represents the number of genes involved, and the size indicates the level of significance for each pathway.

To identify genes whose expressions were regulated by the KDM5 family of proteins, iPSC-CMs were treated with KDM5-C70 (0.5µM) for two weeks and analyzed for gene expression changes by RNA-Seq. The untreated and DMSO-treated cells served as controls. Multidimensional scaling (MDS) plots of the RNA-seq dataset showed a clear separation of the treated and control groups, whereas the two control groups were not distinct from each other (Figure 2D). Likewise, there was no differentially expressed gene (DEGs), defined at a q<0.01 and 1.5-fold change, between the two control groups (untreated vs. DMSO). In contrast, inhibition of KDM5 was associated with the differential expression of 2,415 genes as compared to untreated control cells and 2,427 genes compared to the DMSO-treated cells. A total of 2,372 genes were differentially expressed in the KDM5-C70 iPSC-CMs as compared to DMSO and untreated cells (1,050 upregulated and 1,322 downregulated) (Figure 2E). Heat plots for the DEG genes and clustering between the KDM5-C70-treated and control groups are depicted in Figure 2F. The DEGs were analyzed for the expression of known KDM5A and KDM5B target genes, which showed their increased expression levels in the iPSC-CMs treated with KDM5-C70, in agreement with the known inhibitory role of the KDM5 proteins on gene expression (Figure 2G). Gene set enrichment analysis (GSEA) of the DEGs predicted the upregulation of genes encoding OXPHOS, Fatty acid metabolism, and sarcomere proteins (Figure 2H). The downregulated genes predicted suppression of the expression of genes involved in extracellular matrix formation and inflammation (Figure 2I).

### Identification of H3K4me3 enriched genomic sites in iPSC-CMs

Since KDM5s are histone demethylases that specifically target H3K4me3, genomic loci that are enriched with the H3K4me3 serve as a surrogate for the KDM5 biding sites. To identify the genome-wide H3K4me3 enriched sites, iPSC-CMs treated with the KDM5-C70 and untreated control cells were analyzed by CUT&RUN assay (Cleavage Under Targets & Release Using Nuclease). Approximately, 15 million reads per sample were obtained, which were uniquely mapped to the human genome (GRCH38). Peaks were accessed using the Sparse Enrichment Analysis for CUT&RUN (SEACR) program, which resulted in the identification of a total of 13573 consensus peaks. Heat maps depicting the signal distribution of the H3K4me3 peaks over IgG in the control and KDM5-C70 treated cells are depicted in Figure 3A, which showed a significant signal enrichment in the H3K4me3 antibody group as compared to the background signal in the IgG groups. The heat map of individual samples is shown in Supplementary Figures 3A and 3B. Differential enrichment analysis (q<0.05) identified 4373 up-regulated and 334 down-regulated peaks in iPSC-CMs treated with the KDM5 inhibitor compared to the control group (Figure 3B).

**Figure 3:**
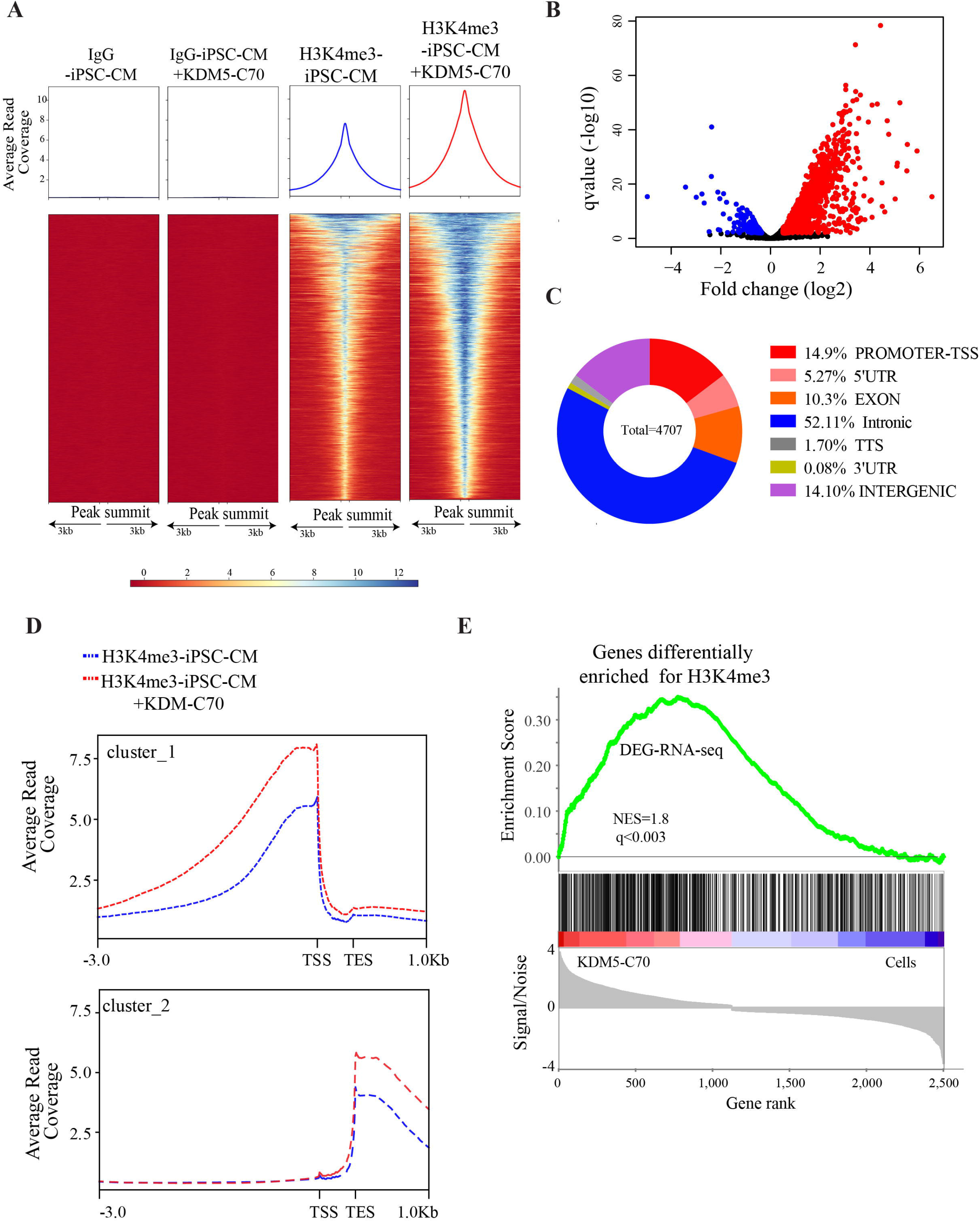
Effect of KDM5 inhibition on genome-wide H3K4me3 distribution. **A)** Heat plot showing cumulative signal intensity for H3K4me3 and IgG in treated and untreated samples. The upper panels show the average profile of the peaks. The lower panels show read density heatmaps around 3Kb from the peak centers. **B)** Volcano plot showing differentially H3K4me3 peaks in the KDM5-C70 treated vs. control groups. Peaks that are differentially up-regulated (higher signal intensity) are shown in red, while down-regulated peaks are shown in blue. **C)** Genome-wide distribution of differentially enriched H3K4me3 peaks. Promoter and upstream regions are defined as the region 3kb upstream and 1kb downstream from the start of transcription. **D)** Distribution of peaks around the transcription start site, gene body and downstream to gene body in control cells (black) and KDM-C70-treated cells (red). **E-F)** GESA plots showing the correlation of RNA levels of differentially expressed genes with the genes that have differentially higher H3K4me3 deposits in KDM5-C70-treated vs. control groups (E) or differential lower H3k4me3 deposits.

The genomic distribution of the H3K4me3 peaks showed marked enrichment at the promoter and TSS upstream regions with a subset of peaks enriched at the intronic cis elements (Figure 3C). In addition to the enrichment of H3K4me3 peaks in the promoter and upstream regions, there was a broader signal distribution of H3K4m3 across the TSS in the KDM5 inhibition group (Figure 3A and 3D). These results imply that inhibition of KDM5 alters the genome-wide distribution of H3K4me3, increased peak intensity (size), and peak width in iPSC-CMs.

To gain insight into the biological significance of these findings, differentially enriched H3K4me3 peaks were analyzed for their effect on gene expression after KDM5 inhibition. Integration of the RNA-seq and the CUT&RUN dataset was performed by analyzing the up-regulated and down-regulated peaks against the DEG. This unbiased analysis showed that of 1126 differentially upregulated genes, 457 had increased H3K4me3 deposits at their promoter and upstream regions after KDM5 inhibition (Figure 3E). The downregulated genes upon treatment with the KDM5 inhibitor did not show reduced deposition of H3K4me3 at their promoter and upstream regions. The data suggest that inhibition of KDM5 led to dysregulation of gene expression in an H3K4me3-dependent manner.

### Inhibition of KDM5 induces expression of genes encoding sarcomere and myofibrillar organization in the hiPSC-CMs

Analysis of the RNA-seq data showed enrichment of the myogenesis-associated genes in iPSC-CMs treated with KDM5-C70 compared to controls. Similarly, GO analysis showed that genes involved in sarcomere and myofibrillar organization were highly enriched in the iPSC-CMs treated with KDM5-C70 (Figure 4A). Furthermore, independent analysis of the transcript levels of several sarcomere genes by qPCR confirmed induction of maturation-specific sarcomere genes TNNI3 (∼ 5-fold), MYL2 (∼ 3-fold), MHY7 (∼ 1.4-fold), TNNT2 (∼1.4-fold), MYOZ2(∼2-fold), TPM1(∼1.6-fold) and ACTC1(∼1.6-fold) as shown in Supplementary Figure 4A. To determine whether changes in gene expression correspond to changes in H3K4me3 levels, selected differentially expressed sarcomere genes were analyzed for the H3K4me3 accumulation at their promoters. Overall, 27 sarcomeric/myogenesis genes showed a concordant increase in their transcript levels and H3K4me3 peaks at their promoters in iPSC-CMs treated with KDM-C70. The corresponding heat map is shown in Figure 4B. Selected examples of H3K4me3 accumulation at the promoter regions of the sarcomere genes implicated in CM maturation^6, 7, 41^ namely, MHY7, MYL2, TNNI3, and MYOM2 are shown in Figure 4C. The data suggest that KDM5 regulates gene expression in iPSC-CMs, including genes involved in sarcomere formation, through genome-wide regulation of H3K4me3 at the target locus.

**Figure 4:**
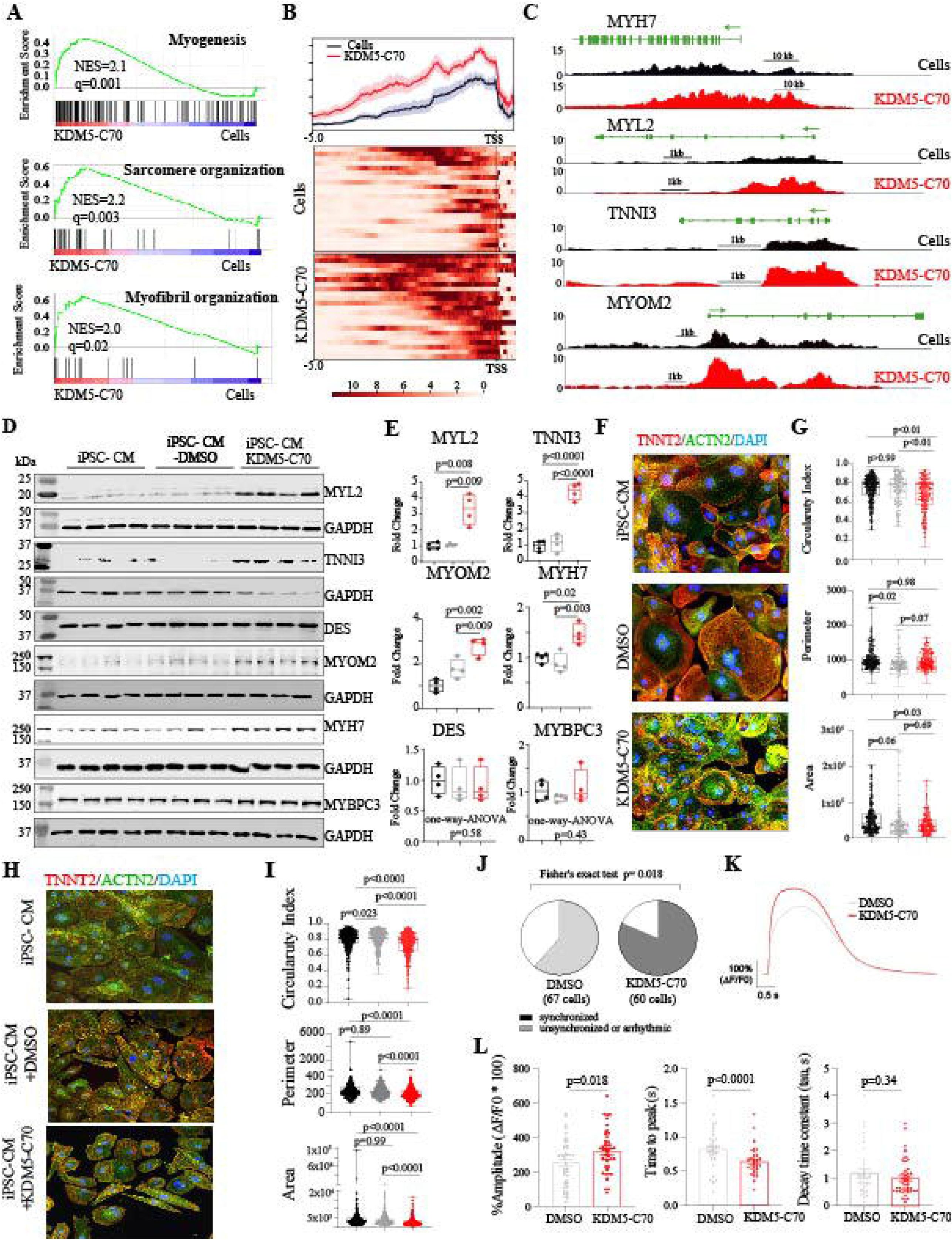
Effect of KDM5 inhibition on sarcomere gene program. **A)** GSEA and GO analysis of DEGs showing induction of the gene program involved in myogenesis, sarcomere formation myofilament formation gene program. **B)** Distribution of H3K4me3 peak density at the TSS of sarcomere genes differentially expressed in KDM5-C70 treated (red) versus control (black) samples and heat plot showing the signal intensity of H3K4me3 in the upstream regions of sarcomere genes in control and KDM5-C70 treated samples. **C)** IGV genome browser trace of H3K4me3 showing peak intensity at the promoter and upstream regions of selected sarcomere genes in control and treated samples. **D-E)** IB analysis showing increased expression of MYL2, TNNI3, MYOM2 and MYH7 and no changes in Desmin and MYBPC3 in KDM5-C70 treated groups. **F)** IF staining with anti-TNNT2 (red), anti-ACTN1 (green) showing sarcomeres and a prominent striated pattern in KDM5C-70 treated cells. Cells were plated on a Matrigel coated glass coverslip. **G)** Calculated circularity index (0 = oblong and 1 = circle), area and perimeter for control, DMSO and KDM5-C70 treated samples. **H-I)** Similar IF staining and quantification on PDMS-coated slides. Pairwise corrected p values are shown in the figure. **J)** Pie chart showing percentage of cells responding to synchronization by pacing in the indicated groups. **K)** Representative traces of intracellular Calcium transient from DMSO (grey) and KDM5-C70 treated cells (red) **L)** Quantitative measurement showing calcium transient amplitude, time to peak and decay constants (Tau) in the DMSO and treated groups.

Protein levels of selected sarcomere genes were determined in the experimental groups by IB, which showed increased expression of MYL2 (4.2±1.1-fold), TNNI3 (4.2±0.4-fold), MYH7 (2.46±0.1-fold) and MYOM2 (1.8±0.3-fold) after KDM5-C70 treatment (Figure 4E and 4F). To quantitatively assess differences in cell morphology of iPSC-CMs upon treatment, we measured cell surface area, cell perimeter, and circularity index. Cells treated with KDM5-C70 showed significantly less circularity, indicating that KDM5-C70 caused the elongation of these cells (Figure 4F and 4G). Cells treated with KDM5-C70 were approximately 30% smaller than the untreated or DMSO-treated cells. A hallmark of cardiomyocyte maturation is the assembly of organized arrays of myofibrils. To visualize the regularity of myofibrillar assembly, iPSC-CMs were immunostained for the sarcomere protein actinin (ACTN2) and troponin T2 (TNNT2). Treatment with KDM5-C70 resulted in regularly arranged sarcomeres, while control iPSC-CMs showed sparse disorganized sarcomeres (Figure 4F). Since the extracellular matrix (ECM) plays an important role in iPSC-CM maturation, we plated the iPSC-CMs on another ECM prepared by PDMS/Matrigel coating of glass coverslips ^42, 43^. The results were remarkable for an increased number of elongated cells with organized sarcomeres in the KDM5-C70-treated groups compared to the untreated cells on the same substrate (Figure 4H and Figure 4I).

To determine the effects of KDM5 inhibition on cardiomyocyte excitation-contraction coupling, we measured Ca^2+^ transients elicited by electrical field stimulation. iPSC-CMs (after 15 days of treatment with vehicle or KDM5-C70) were plated at 50% confluency, loaded with Fluo-4 AM, and their fluorescence intensity was measured using IonOptix. Cells were continuously paced at room temperature at a frequency of 0.2 Hz to achieve steady-state Ca^2+^ levels. KDM5-C70 treatment significantly increases synchronization to pacing as compared to those treated with the vehicle (Figure 4J). Interestingly, synchronized KDM5-C70-treated cardiomyocytes exhibited an increased Ca^2+^ transient amplitude with faster kinetics compared to those observed treated with a vehicle (Figure 4K-L). Together, this evidence suggests that inhibition of KDM5 improves iPSC-CM excitation-contraction coupling.

### Inhibition of KDM5 enhances oxidative phosphorylation and fatty acid beta-oxidation in hiPSC-CMs

The prenatal heart relies on glucose and lactate for energy, but in the immediate postpartum period when dietary fat is abundant, the heart switches to fatty acid oxidation (FAO) as the primary energy source^6, 7, 44^. Consistent with the metabolic switch during maturation of cardiomyocytes, KDM5-C70 treatment induced the expression of several genes involved in fatty acid metabolism (Figure 5A). Fatty acid metabolism consists of several steps, including FA transport within the mitochondria, oxidation to produce Acyl-CoA for the TCA cycle, and finally the generation of ATP by OXPHOS. RNA-seq analysis revealed that genes involved in different stages of FA metabolism were induced by KDM5-C70 treatment. QPCR analysis for the transcripts of genes involved in fatty acid transport CPT1A, CPT1B, and SLC27A showed a 1.5- to 3-fold increase upon KDM5 inhibition. Likewise, genes involved in acyl-CoA degradation via oxidation, such as ACADVL, HADHA, and HADHB, were induced by KDM5-C70 treatment (Figure 5B and Supplementary Figure 5B). The H3K4me3 signal for these specific genes showed a predominant accumulation at their promoter regions in the cells treated with the KDM5 inhibitor (Fig. 5C). Figure 5D shows a selection of peaks demonstrating both increased size and width at the promoters of the CPT1B, MLYCD, and ACADS genes. Finally, the protein levels of these genes were assessed, which showed elevated levels for CPT1B (2.09±0.3), ACADVL (7-fold), ACADM (2.12±0.1), and ECHDC3 (13.3±2.2) as shown in Figure 5E and 5F.

**Figure 5:**
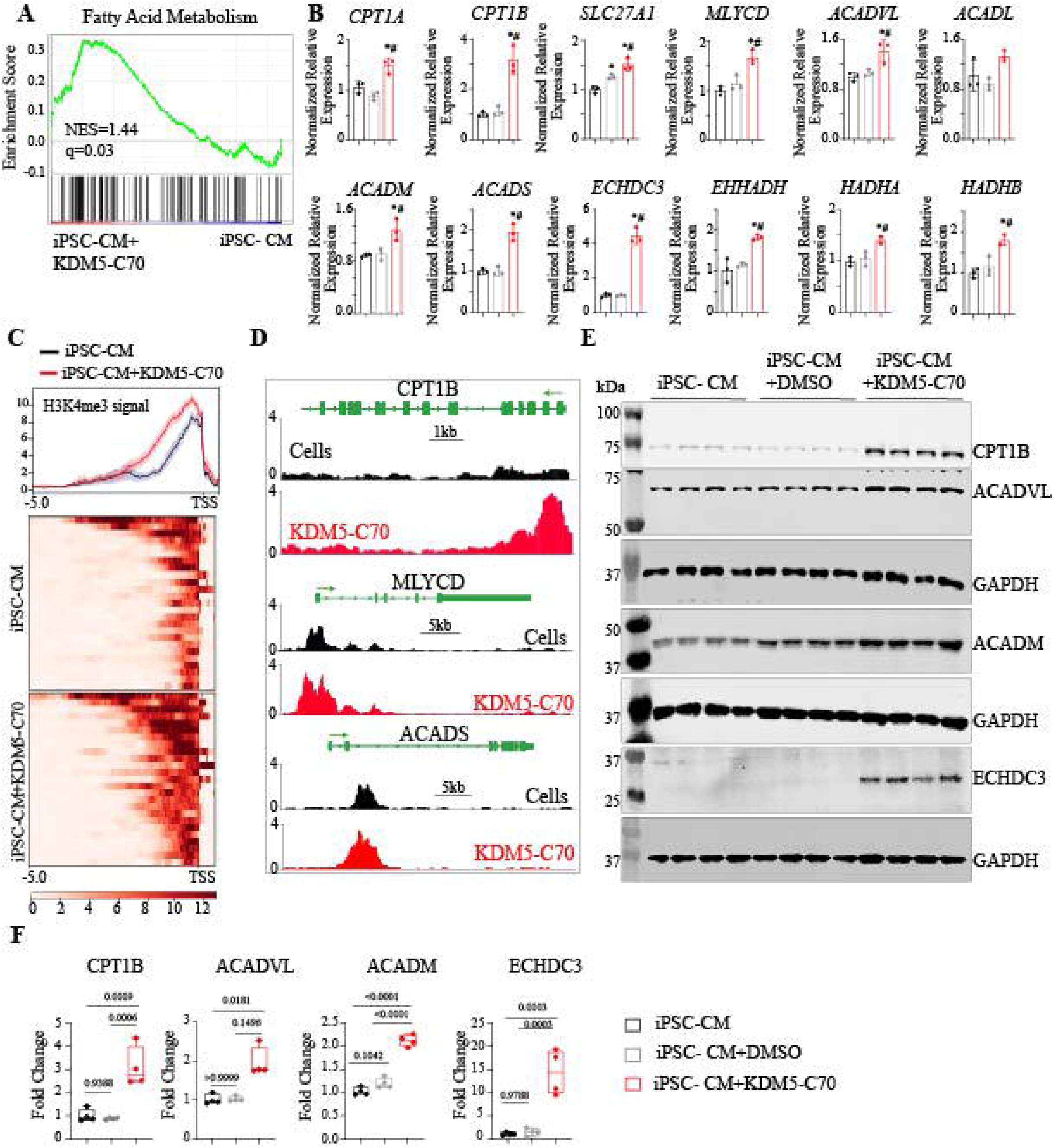
KDM5 inhibition leads to induction of FAO gene program. **A)** GSEA showing activation of fatty acid metabolism gene signature. **B)** QPCRs of genes involved in FA transport and FA oxidation demonstrate induction in KDM5-C70 treated cells. **C)** Heat plots showing distribution profile of peak density and peak width enrichment near the TSS of fatty acid metabolism genes in cells and KDM5-C70 treated samples **D)** IGV genome browser traces of H3K4me3 on selected genes involved in fatty acid metabolism in the control (black) and KDM5-C70 treated samples (red). **E)** IB analysis showing increased expression of the fatty acid transporter CPT1B, and the enzymes involved in beta-oxidation of fatty acids namely ACADVL (Krushkal-Wallis test p =0.0048), ACADM and ECHDC3 in treated cells. **F)** Quantitative data showing the fold change in the expression of proteins as shown in E. Pairwise corrected pvalues are shown in the figure.

Transcripts of genes involved in OXPHOS were increased in iPSC-CMs treated with KDM5-C70, as evident from the GESA plots shown in Figure 6A and the QPCR data shown in Figure 6B. However, the CUT-and-RUN assay showed accumulation of the H3K4me3 in only a few genes (6/80) in the treatment group, including the transcription factor ESRRA, which is a major regulator of OXPHOS gene expression in cardiomyocytes (Figure 6C) ^45-47^. QPCR analysis confirmed the induced expression of ESRRA (2.1 ±0.2) in the KDM5-C70-treated group as shown in Figure 6D. The ESRRA transcript was also increased in another iPSC-CM line treated with KDM5-C70 (Supplementary Figure 5C). Consistent with changes in the RNA levels, protein levels of ESRRA were induced by 2.4±0.36-fold after KDM5-C70 treatment as shown in Figure 6, panels E and F. Finally, transcript levels of ESRRA target genes, which are mainly involved in OXPHOS, were induced by KDM5-C70 treatment (Figure 6G). However, mitochondrial DNA content, which reflects an increase in mitochondrial number, was not increased by KDM5-C70 treatment (Supplementary Figure 5A). Overall, our data suggest that inhibition of KDM5 induces ESRRA and thereby contributes to transcriptional induction of OXPHOS gene expression.

**Figure 6:**
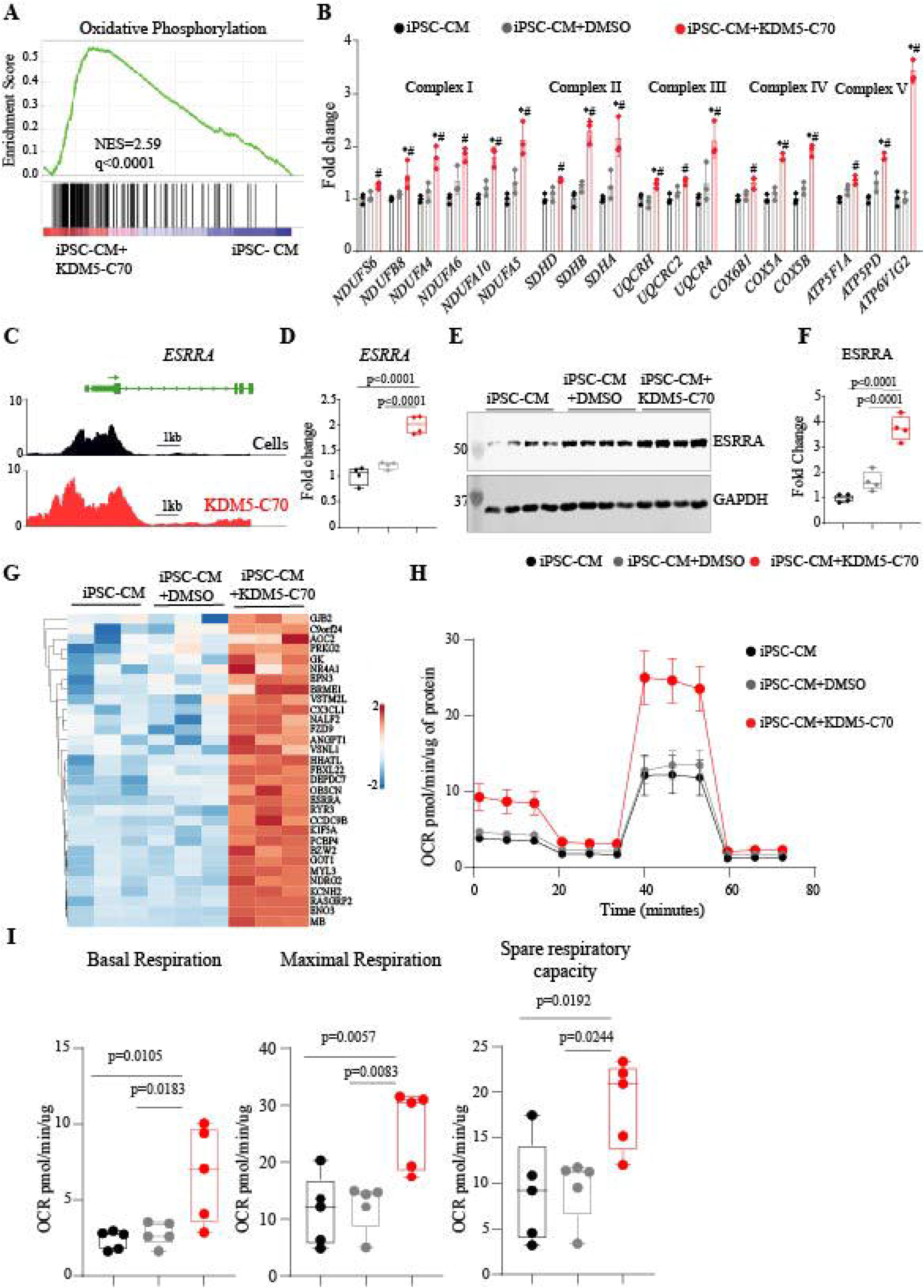
Effect of KDM5 inhibition on OXPHOS. **A)** GSEA from DEGs-predicted activation of gene signature of OXPHOS upon KDM5 inhibition. **B)** QPCR data showing transcript levels of selected OXPHOS genes from complex I to V. **C)** IGV genome browser traces of H3K4me3 around TSS of *ESRRA* gene **D)** QPCR data showing transcript levels of *ESRRA*. **E and F)** IB showing induction of ESRRA protein after treatment with KDM5-C70**. G)** Heat plots showing targets of ESRRA as induced in the treated groups. **H)** Panel shows kinetic data for Real-time oxygen consumption rate (OCR) measurements of hiPSC-CMs by Seahorse extracellular flux analyzer in the controls and treated groups. Average data from five independent experiments each constituting an average 10 wells/ experiment is shown. Error bar indicates SEM. **I)** KDM5-C70 treatment demonstrated higher respiratory rate under baseline conditions and after mitochondrial decoupling. Quantitative data obtained from five independent experiments each constituting an average of 10 wells is shown. Pairwise corrected p-values after One way ANOVA are shown

Finally, to determine the effect of gene expression changes on OXPHOS, an extracellular flux assay was performed to measure real-time oxygen consumption rate (OCR) (Figure 6H). Inhibition of KDM5 in the iPSC-CMs increased the basal respiratory rate. In addition, there was a 2-fold difference in maximal respiration and spare capacity, between the KDM5-C70-treated and untreated cells (Figure 6I), demonstrating improved aerobic respiration and ATP production in KDM5-C70 treated cells. Taken together, these results support the role of KDM5 in iPSC-CM metabolic maturation through FAO and OXPHOS regulation.

## Discussion

The results demonstrate that inhibition of KDM5 activity by a pan-KDM5 small molecule inhibitor KDM5-C70 is sufficient to induce H3K4 methylation, increase H3K4me3 level, and activate gene expression. Inhibition of KDM5 in immature iPSC-derived cardiomyocytes resulted in a gene expression reprogramming that resembles that of mature CM. Specifically, inhibition of KDM5 led to the induction of genes involved in fatty acid oxidation and OXPHOS, leading to improved FAO and OCR in iPSC-CMs. Similarly, genes encoding sarcomere proteins were induced by KDM5 inhibition, resulting in the striated organization of the sarcomere. Mechanistically, the data suggest that KDM5 inhibition shifted the iPSC-CM toward maturation through A) direct regulation of genes involved in FAO, OXPHOS, and sarcomere formation and B) regulation of the transcription factors ESRRA, a known regulator of cardiomyocytes maturation ^46, 47^. Our results are consistent with recent reports showing the role of ESRRA in cardiomyocyte maturation mainly through the regulation of genes involved in OXPHOS and FAO and creating of an epigenetic link to the maturation gene program regulated by ESRRA. It has been proposed that KDM5A both induces and represses mitochondrial biogenesis and function, however, its role in regulating the energy metabolism of cardiomyocytes is unknown. Recently, we showed that in a mouse model of laminopathies, KDM5A was induced and associated with the repression of its target genes involved in OXPHOS^16^. Our current results indicate that the KDM5 family of proteins can regulate energy metabolism in cardiac myocytes through multiple target genes that not only affect mitochondrial gene expression but are also involved in fatty acid uptake and oxidation.

Our studies also point to the role of KDM5 in the regulation of the expression of sarcomere proteins. Inhibition of KDM5s resulted in increased deposition of H3K4me3 on the promoter regions of several sarcomere genes, namely MYL2, TNNI3, and MYH7 among others, and led to an increase in their RNA and proteins levels. These expression changes are conserved and vital for the transition from prenatal to postnatal cardiomyocytes and are therefore an integral part of the maturation process of cardiomyocytes in adults. The results are intriguing and unexpected and will open new avenues for targeting the sarcomere gene expression by modulating KDM5 activities upon further investigation and validation in the *in vivo* models.

The molecular basis of KDM5-mediated gene regulation in cardiomyocytes remains unknown. Our data indicate that KDM5s are one of the major regulators of H3K4me3 demethylation in iPSC-CMs. Likewise, our data points to a direct role of H3K4me3 in gene regulation as the changes in the H3K4me3 levels after KDM5-C70 treatment strongly correlate with changes in gene expression. The precise role of H3K4me3 in gene expression might be context-dependent, but recent data have shown that H3K4me3 regulates promoter-proximal pause release of RNA polymerase II ^48^.

Our data indicate that KDM5 inhibition reprograms gene expression towards maturation of iPSC-CM however complete maturation of these cells may require additional optimized substrates and stimuli to fully reprogram these cells to a mature state. Finally, the individual contribution of each KDM5 to cardiac gene regulation remained unknown. Nonetheless, our data provide a first insight into the role of KDM5s in cardiomyocytes and provide evidence that inhibition of KDM5 has a notable effect on the sarcomere, OXPHOS, and FAO genes that prime the iPSC-CMs towards maturation.

## Conclusion

Overall, our data confirm the importance of KDM5 in regulating cardiomyocyte maturation and demonstrate its effect on OXPHOS and sarcomere gene regulation. Remarkably, KDM5 on the one hand represses CM maturation and consequently reduces the expression of ESRRA and on the other hand targets genes involved in sarcomere formation. Therefore, we conclude that KDM5 represents a novel target in the regulation of cardiac myocyte maturation and may have potent therapeutic effects in combination with other known myocyte maturation strategies.

## Acknowledgements

We thank the MDACC Epigenomics Profiling Core Facility for their assistance with the CUT&RUN assays.

