## Supplementary Figures for "Histone demethylase KDM5 regulates cardiomyocyte maturation by promoting fatty acid oxidation, oxidative phosphorylation, and myofibrillar organization"

**Supplementary Figure 1**

**A)** Human-induced pluripotent stem cells were differentiated into CM and RNA was collected at the indicated time points. qPCR analysis for mesodermal markers T (Brachyury) and MESP1 are shown. **B)** IF staining using anti-TNNT2 antibody (red) showing that the differentiation protocol produced 85.34 20.8% TNNT2+ cells. **C)** q-PCR analysis showing the expression of KDM5D along with KDM5A, KDM5B and KDM5C in the CHIP22 cell line.

**Supplementary Figure 2:**

IB analysis for H3K27me3 and H3K9me3 after treatment with different concentrations of KDM5-C70. The values at the bottom of the blots represent the change in fold for each group.

**Supplementary Figure 3**

Heat plots and peak profile plots showing the read density for all 4 groups used in the CUT&RUN assay for H3K4me3 normalized to IgG for control and KDM5-C70 treated cells.

**Supplementary Figure 4:**

**A)** Normalized relative expression of sarcomeric genes in control, DMSO and KDM5-C70 treated groups as analyzed by q-PCR. **B)** Normalized relative expression of sarcomeric genes validated in an independent cell line CHIP22S upon KDM5-C70 treatment. **C)** Normalized relative expression of genes involved in Calcium handling in cardiomyocytes.

**Supplementary Figure 5: A)** Normalized relative expression of genes involved in fatty acid metabolism and oxidative phosphorylation after treatment with KDM5-C70 in an independent cell line CHIP22S corresponding to main Figure 5B and 6B. **B)** Normalized relative expression of transcription factor ESRRA in CHIP22S cell line after KDM5-C70 treatment. **C)** qPCR analysis of ratio of Mitochondrial DNA (ND1) to nuclear DNA (Beta globin and 18SrRNA)

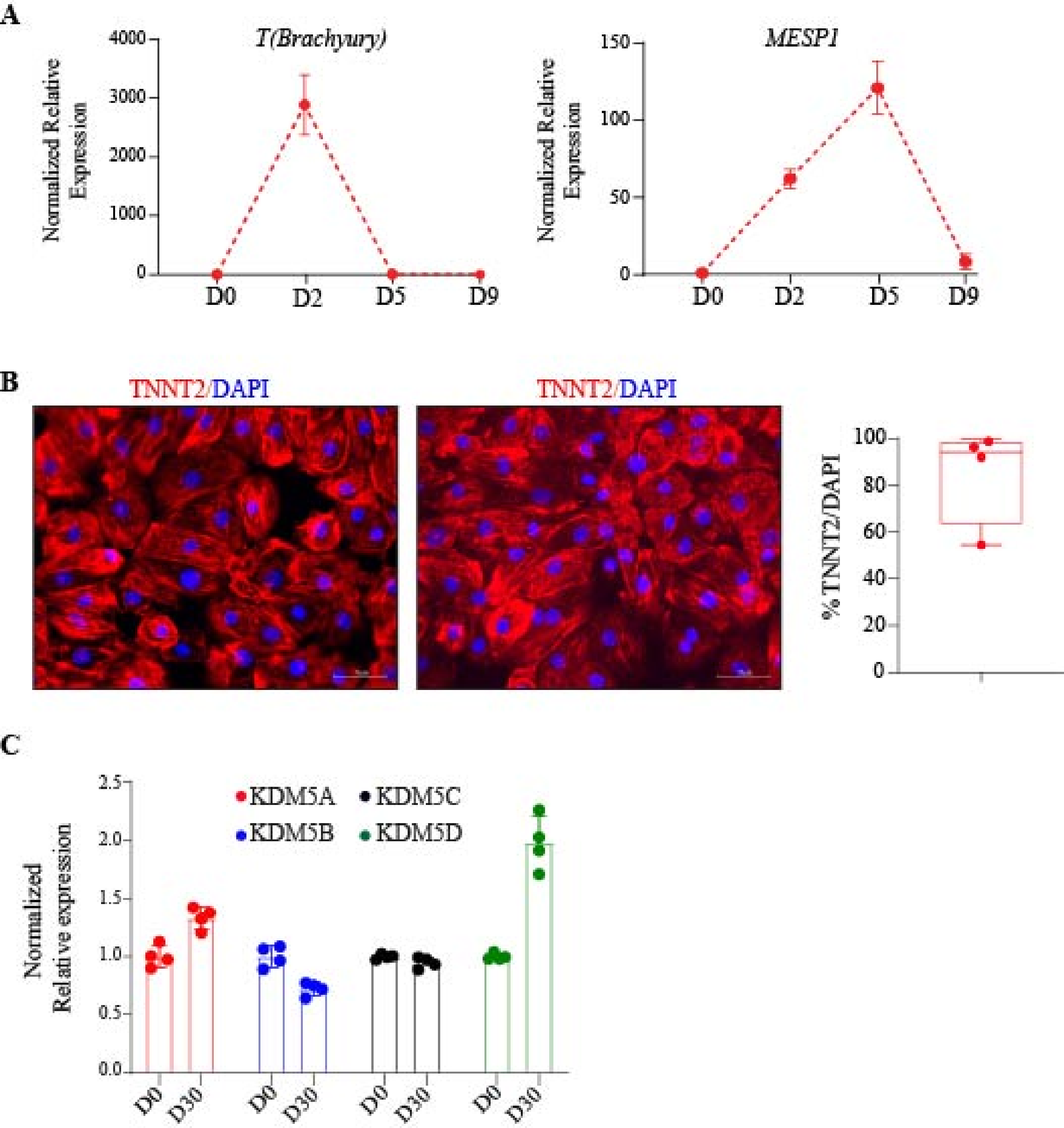

A
T(Brachyury)
Normalized Relative Expression
4000
3000
2000
1000
0
D0
D2
D5
D9
MESP1
150
100
50
B
TNNT2/DAPI
TNNT2/DAPI
% TNNT2/DAPI
100
80
60
40
20
C
KDM5A
KDM5C
KDM5B
KDM5D
Normalized Relative expression
2.5
2.0
1.5
1.0
0.5
0.0
D30

A

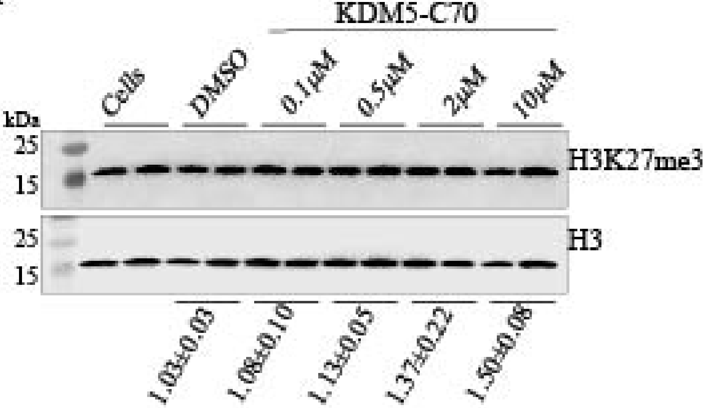
KDM5-C70
Cells
DMSO
0.1μM
0.5μM
2μM
10μM
kDa
25
15
H3K27me3
25
15
H3
1.03±0.03
1.08±0.10
1.13±0.05
1.37±0.22
1.50±0.08

B

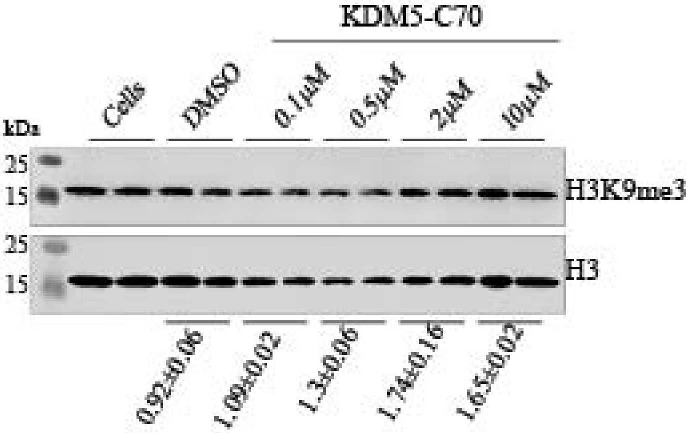
KDM5-C70
Cells
DMSO
0.1μM
0.5μM
2μM
10μM
kDa
25
15
H3K9me3
25
15
H3
0.92±0.06
1.09±0.02
1.3±0.06
1.74±0.16
1.65±0.02

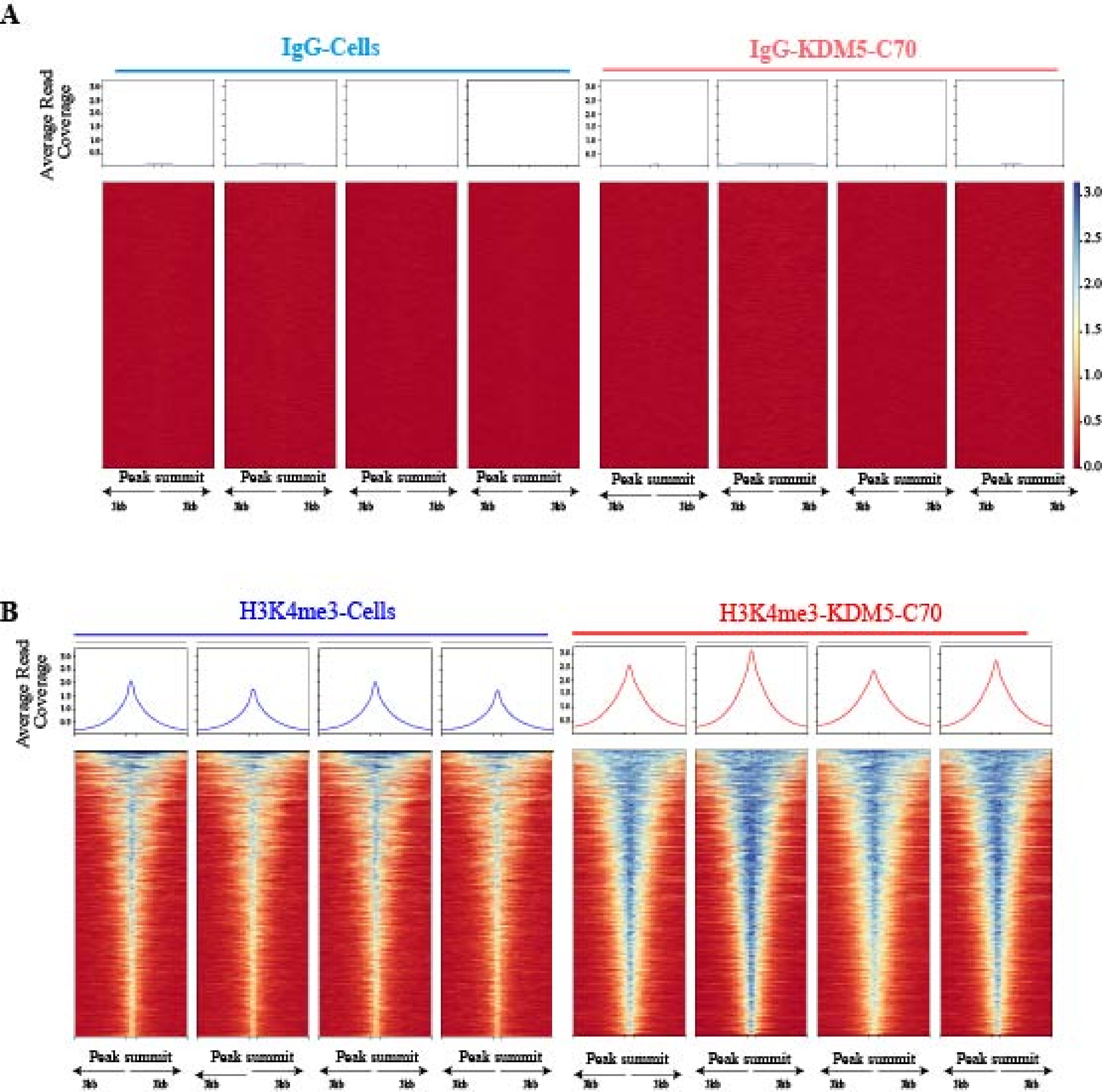
A
IgG-Cells
IgG-KDM5-C70
Average Read Coverage
Peak summit
3.0
2.5
2.0
1.5
1.0
0.5
0.0
B
H3K4me3-Cells
H3K4me3-KDM5-C70
Average Read Coverage
Peak summit

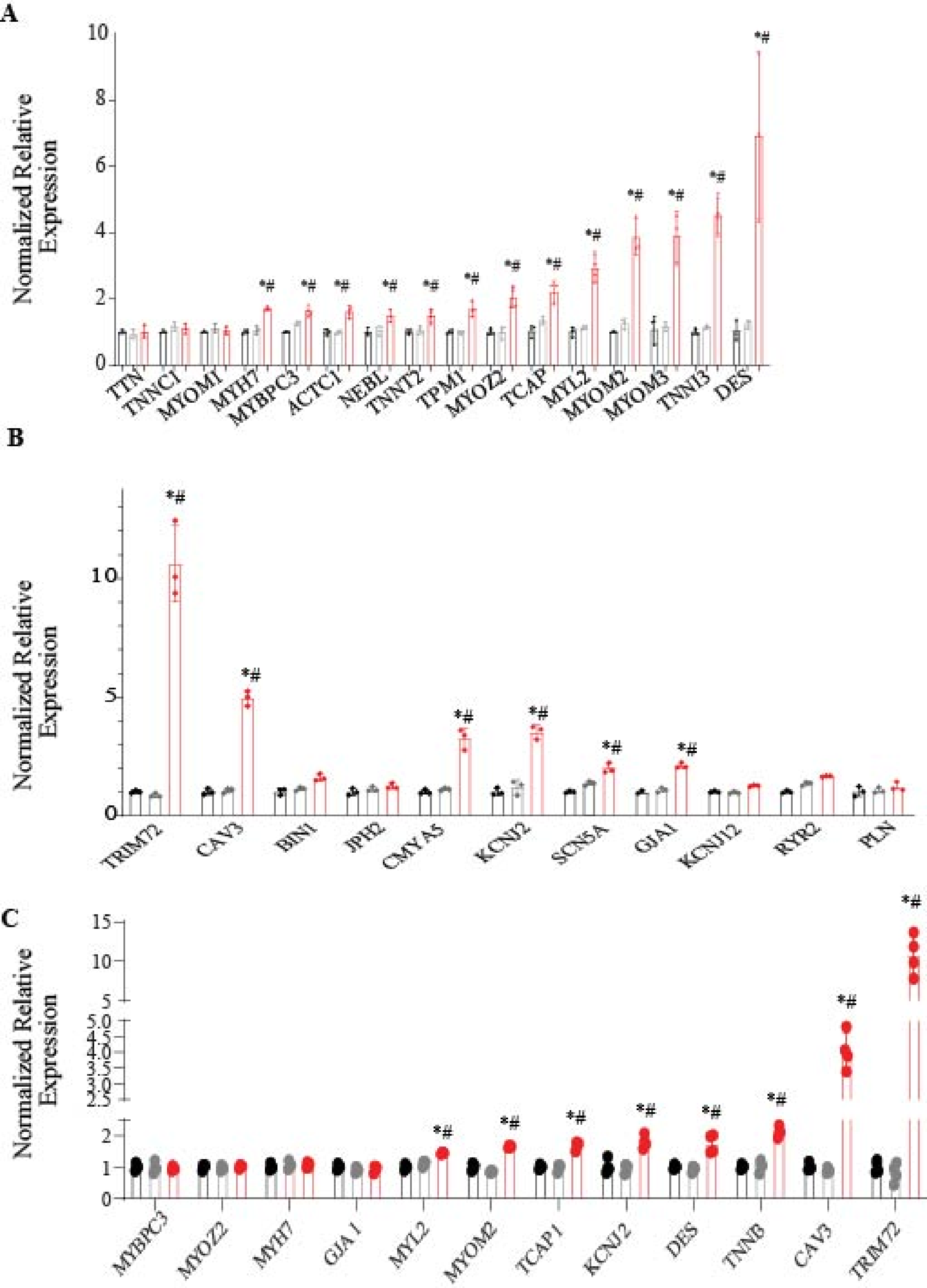
A
Normalized Relative Expression
0
2
4
6
8
10
*#
TTN
TNNC1
MYOM1
MYH7
MYBPC3
ACTC1
NEBL
TNNT2
TPM1
MYOZ2
TCAP
MYL2
MYOM2
MYOM3
TNNI3
DES
B
Normalized Relative Expression
0
5
10
*#
TRIM72
CAV3
BIN1
JPH2
CMYA5
KCNJ2
SCN5A
GJA1
KCNJ12
RYR2
PLN
C
Normalized Relative Expression
0
1
2
5
10
15
*#
MYBPC3
MYOZ2
MYH7
GJA 1
MYL2
MYOM2
TCAP1
KCNJ2
DES
TNNI3
CAV3
TRIM72

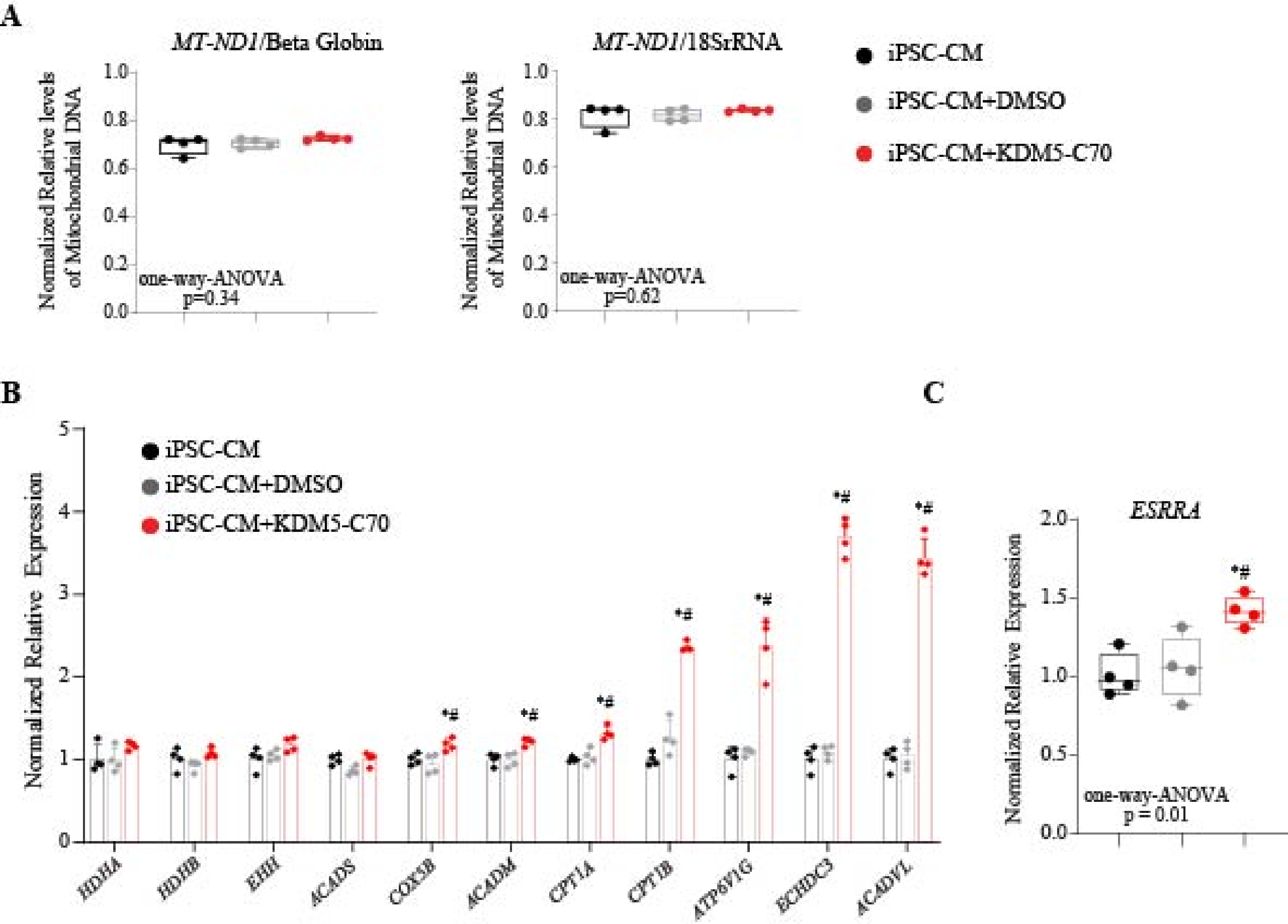
A
MT-ND1/Beta Globin
MT-ND1/18SrRNA
Normalized Relative levels of Mitochondrial DNA
one-way-ANOVA
p=0.34
one-way-ANOVA
p=0.62
iPSC-CM
iPSC-CM+DMSO
iPSC-CM+KDM5-C70
B
Normalized Relative Expression
HDHA
HDHB
EHH
ACADS
COX5B
ACADM
CPT1A
CPT1B
ATP6V1G
ECHDC3
ACADVL
C
ESRRA
one-way-ANOVA
p = 0.01
